# Changes in social groups across reintroductions and effects on post-release survival

**DOI:** 10.1101/430280

**Authors:** Victoria R. Franks, Caitlin E. Andrews, John G. Ewen, Mhairi McCready, Kevin A. Parker, Rose Thorogood

## Abstract

Reintroductions are essential to many conservation programmes, and thus much research has focussed on understanding what determines the success of these translocation interventions. However, while reintroductions disrupt both the abiotic and social environments, there has been less focus on the consequences of social disruption. Therefore, here we investigate if moving familiar social groups may help animals (particularly naïve juveniles) adjust to their new environment and increase the chances of population establishment. We used social network analysis to study changes in group composition and individual sociality across a reintroduction of 40 juvenile hihi (*Notiomystis cincta*), a threatened New Zealand passerine. We collected observations of groups before a translocation to explore whether social behaviour before the reintroduction predicted associations after, and whether reintroduction influenced individual sociality (degree). We also assessed whether grouping familiar birds during temporary captivity in aviaries maintained group structure and individual sociality, compared to our normal translocation method (aviaries of random familiarity). Following release, we measured if survival depended on how individual sociality had changed. By comparing these analyses with birds that remained at the source site, we found that translocation lead to re-assortment of groups: non-translocated birds maintained their groups, but translocated juveniles formed groups with both familiar and unfamiliar birds. Aviary holding did not improve group cohesion; instead, juveniles were less likely to associate with aviary-mates. Finally, we found that translocated juveniles that lost the most associates experienced a small but significant tendency for higher mortality. This suggests sociality loss may have represented a disruption that affected their ability to adapt to a new site.

## Introduction

Reintroduction, returning species to parts of their range where they have become extinct (IUCN/SSC, 2013), is important for many conservation programmes (Armstrong and Seddon, 2008). The process of moving animals to a new site (“translocation” (IUCN/SSC, 2013)) and overcoming post-release effects during “establishment” (IUCN/SSC, 2013) are critical to the success of reintroductions (Fischer and Lindenmayer, 2000; Bennett *et al*., 2012; Parker *et al*., 2012; Miskelly and Powlesland, 2013; Armstrong *et al*., 2017). Novelty of the post-release environment appears to be a major challenge to survival because animals need to avoid starvation and predation with little personal experience of the release site (Letty *et al*., 2003; Pinter-Wollman, Isbell and Hart, 2009; Batson, Abbott and Richardson, 2015). Thus, key remaining questions in reintroduction biology centre around how animals adjust successfully to their new environment and what can increase their post-release survival (Anthony and Blumstein, 2000; Armstrong and Seddon, 2008).

Reintroductions change the abiotic environment, but also the social environment when the founding group of animals represents a subsample of a larger original population (Ewen, Armstrong, Parker, *et al*., 2012). The composition of groups may be important for establishment (Anthony and Blumstein, 2000; Clarke, Boulton and Clarke, 2003; Armstrong and Seddon, 2008; IUCN/SSC, 2013) because it can affect how animals adjust behaviour as a first-response mechanism to a new environment (Wong and Candolin, 2014). Animals may prefer to associate with and learn from familiar peers when finding food or avoiding predation (Atton & Galef, 2014; Lachlan, Crooks, & Laland, 1998; Schwab, Bugnyar, Schloegl, & Kotrschal, 2008; but see Ramakers, Dechmann, Page, & O’Mara, 2016) and in novel environments, collective group knowledge may become more important because it can offset each individuals’ limited personal experience (King and Cowlishaw, 2007; Pinter-Wollman, Isbell and Hart, 2009). Associations can also affect the likelihood that animals disperse post-release, as more cohesive groups are less likely to split up (Blumstein, Wey and Tang, 2009; Snijders *et al*., 2017). Therefore, if animals lose previous social connections during reintroductions there may be consequences for population stability, for both the translocated population as well as the remaining source population (Blanchet, Clobert and Danchin, 2010; Pinter-Wollman *et al*., 2013).

To understand how group structure and familiarity impacts on translocation success, we therefore first need to determine if groups remain together when they are moved to a new site. One challenge in wild animal groups is there may be limited knowledge of familiarity before translocation. For example, studies in New Zealand bird species (tīeke/saddleback, *Philesturnus carunculatus rufusater*; toutouwai/North Island robin, *Petroica longipes*) and howler monkeys (*Alouatta seniculus*) found that pre-capture familiarity was not maintained over translocation (Armstrong, 1995; Armstrong and Craig, 1995; Richard-Hansen, Vié and De Thoisy, 2000). However, these species are territorial, and the studies also defined familiarity from short-term binary measures (individuals in the same place upon capture were “familiar”, versus “non-familiar”). When longer-term measures of familiarity have been used for more social groups (such as families or colonies) there is evidence that group composition remains similar before and after reintroduction (Clarke, Boulton and Clarke, 2003; Shier, 2006; Pinter-Wollman, Isbell and Hart, 2009) and that maintaining groups results in higher post-release survival (Shier, 2006). Therefore, capturing group familiarity over a longer time period for social species may be required to assess the importance of maintaining or disrupting relationships over translocations.

Along with the identity of members of a social group, an animal’s number of social connections may also affect how well it adjusts in a novel environment. Some individuals have many associates while others have few (Krause, Lusseau and James, 2009); generally, more social animals acquire new behaviours more quickly, likely because they have more potential sources of information to use (Aplin *et al*., 2012; Snijders *et al*., 2014). In a new and unknown environment (such as a release site), where animals need to acquire new information (for example, to discover new foraging sites), having many social connections may therefore be beneficial. Number of social connections may be especially important if animals are particularly reliant on learning with others, for example juveniles who need to overcome their own limited personal experience (Letty, Marchandeau and Aubineau, 2007; Nuñez, Adelman and Rubenstein, 2015). However, by disrupting the social environment, reintroductions may change an individual’s associations. Little is yet known about individual-level consistency in sociality, and there is limited research exploring the consequences of extensive social disruption (Nuñez, Adelman and Rubenstein, 2015; Firth *et al*., 2017). However, evidence from one study suggests that animals (juvenile feral horses *Equus caballus*) with more social connections may better survive changes in groups such as loss of members, compared to less social peers (Nuñez, Adelman and Rubenstein, 2015); this may be because they lose proportionally fewer associates. Individual sociality may therefore also be important for population stability during establishment (Modlmeier *et al*., 2014; Snijders *et al*., 2017).

Familiarity may also develop between previously unfamiliar individuals if animals are held in temporary captivity for disease screening or to acclimate them to the release site (IUCN/SSC, 2013; Batson, Abbott and Richardson, 2015). This has implications for both group structure and individual sociality, if it promotes cohesion and increases associations. In some social species such as African wild dogs (*Lycaon pictus*) and lions (*Panthera leo*), groups formed during pre-release integration were more likely to remain together post-release, while translocations with non-integrated individuals failed (Gusset, Slotow and Somers, 2006; Hunter *et al*., 2007). However, there also are cases where temporary captivity did not significantly improve group cohesion over immediately-released groups (Clarke, Boulton and Clarke, 2003), or groups disbanded even if translocated together (Fritts, Paul and Mech, 1984). Further, there are other implications of delayed release and in some species it can lead to reduced post-release survival (Castro *et al*., 1994; Richardson *et al*., 2013) or increased stress (Batson *et al*., 2017). Thus, the benefits of temporary captivity are likely species-specific (Moseby, Hill and Lavery, 2014) and a clear understanding of a variety of different advantages and disadvantages of this strategy (including social cohesion) are important to evaluate its use for the species in question (IUCN/SSC, 2013).

Social network analysis provides a way to examine the detailed relative changes in group composition and individual social traits, but as yet has had limited application to studies of conservation value (Wey *et al*., 2008; Snijders *et al*., 2017). By collecting repeated observations of co-occurring individuals, we can determine relative familiarity across a population and define “communities” of frequently co-occurring individuals (Krause *et al*., 2015). We can also calculate individual-level metrics, such as number of associates (“degree centrality”) (Krause *et al*., 2015). Following translocation, we can assess changes in network structure and individual metrics (Snijders *et al*., 2017). If we need to identify particular social characteristics of groups or individuals that are beneficial to reintroductions, social network analysis could inform conservation practice. Further, using translocation as an experimental platform can help us test how group- and individual-level network changes impact on population stability, including survival, information flow and disease dynamics (Snijders *et al*., 2017). As such, opportunities where we can implement social network analysis when testing effects of changing social groups in species of conservation concern provide valuable examples that both inform conservation practice, and help understand the broader ecological and evolutionary consequences of social networks (Pinter-Wollman *et al*., 2013; Kurvers *et al*., 2014; Formica *et al*., 2016; Firth *et al*., 2017).

Here, we use a translocation of hihi (stitchbird, *Notiomystis cincta*) to test fitness effects of network structure, and assess whether maintaining sociality can improve the outcome of a translocation. This species is a threatened New Zealand passerine (Birdlife International, 2017) which was once widespread across the North Island. Following the introduction of non-native predators when humans arrived in New Zealand, hihi became restricted to a single off-shore island (Hauturu-o-Toi/Little Barrier Island). Since the 1980s a major aim for conservation of this species has been to establish re-introduced populations in predator-controlled areas, and the most recent hihi translocations have involved moving juvenile birds. This cohort appears to be particularly social: juveniles form groups for several months at the end of the breeding season and interact, for example with “play”-like behaviour and allopreening. However, it is unknown whether translocation alters these social groups or what the consequences may be for establishment of populations. We used the opportunity of a translocation in 2017 to test our predictions that: (1) translocated hihi will group with more familiar individuals from either before the translocation, or based on who they were held with during temporary captivity; (2) individuals will remain consistent in their sociality before and after translocation; and (3) any changes in social behaviour will affect survival after translocation.

## Methods

### SOURCE AND RELEASE SITE

In 2017 we reintroduced hihi to Rotokare Scenic Reserve (“release site”, 39°27’15.4”S 174°24’33.0”E) from Tiritiri Matangi Island (“source site”, 36°36’00.7”S 174°53’21.7”E). The source site is a 220ha island scientific reserve of replanted and remnant native fauna which is free of non-native mammalian predators. Hihi were reintroduced to the island in 1995 (Armstrong and Ewen, 2001), and the population (numbering c. 270 in 2017) is now the main source of birds for ongoing translocations to other sites. The release site (230ha, including a 17.8ha lake) is a mainland site of old-growth native forest surrounded by a fence that excludes non-native mammalian predators. Hihi had been locally extinct at this site and in the surrounding region for c.130 years prior to the reintroduction (Angher, 1984).

### DEFINING FAMILIARITY BEFORE TRANSLOCATION

Between 17^th^ January – 19^th^ March we collected 229 hours of observational surveys of 105 individuals to determine familiarity at the source site before translocation. To observe as many juveniles as possible we carried out surveys in nine forested gullies (including the three main group sites occupied by juveniles that year, results not presented here) and at six permanent supplementary feeding stations on the island. This ensured we observed associations among juveniles commonly seen at group sites and also associations with the few juveniles that did not frequent these sites (17/108 juveniles were never seen at group sites). During each one-hour survey we recorded the identities of all juveniles seen within a 10-metre radius of the observer (VF). All hihi have an individual combination of coloured leg rings (applied to nestlings during routine nest monitoring) so each could be identified by sight. We assigned juveniles to the geographical location where they were observed: 40 birds were only ever recorded in the northernmost groups (“north”), 16 at the southern end of the island (“south”) and the remaining 49 mixed between the two (mixed).

Next, we constructed a “group-by-individual” (GBI) matrix where a group comprised any juveniles seen within 15 minutes of the preceding bird. If we did not see any birds during this time, we considered the next juveniles encountered to be part of a new group. This “gambit of the group” approach (Whitehead, 2008) was based on previous observations and analysis of hihi social behaviour, where the majority of groups (and individuals) were recorded in an area for a maximum of 15 minutes (Appendix 2.1). Using the GBI, we built a weighted association network in R (version 3.5.0) (R Core Team, 2017) using the “get_network” function in the R package asnipe (version 1.1.9) (Farine, 2013). Weighted networks provided a more detailed measure of familiarity rather than binary familiar/unfamiliar: each “edge” connecting two juveniles represented at least one co-occurrence in a group, so repeated co-occurrences (and stronger edge weights) would indicate that juveniles were more familiar. We detected

“communities” of frequently co-occurring individuals in the network using the community detection algorithm of Clauset et al. (2004) implemented with the “fastgreedy.community” function (igraph R package version 1.0.9, (Csárdi and Nepusz, 2006)). We ensured that assigned communities were robust following the method of Shizuka & Farine (2016): we generated bootstrapped replicates of the observed network by resampling observations of groups before translocation, and in each bootstrapped network we calculated assortment by the community assigned to each juvenile from the observed network. This allowed us to determine if the observed community structure was robust compared to random expectation by calculating the metric *r_comm_*. If *r_comm_* = 1, all replicated networks result in the same community structure as the observed network; conversely, *r_comm_* = 0 means that assignments are random compared to original assigned communities (*r_comm_* > 0.5 is considered “robust” (Shizuka and Farine, 2016)). We assigned each juvenile a number (1–6) corresponding to its network community.

### TRANSLOCATION

On 27^th^ – 28^th^ March, 40 hihi were caught in mist nets or at feeding stations at the source site using a standard catching technique for this population. After capture, each bird was transported individually to be processed immediately for disease screening (Ewen, Armstrong, Empson, *et al*., 2012). After processing, each bird was released into one of three pre-existing aviaries which have been used in many translocations from the source site (each measuring approximately 5×3×2.5 metres). The aviaries were one large enclosure divided into three flights and filled with dense natural vegetation that limited visual contact between aviaries (aviaries were therefore not in auditory isolation from each other or free-living birds). Each juvenile was assigned to an aviary based on its community in the network before translocation: one aviary contained birds from one community only (“familiar” group), while the remaining two aviaries contained birds from all communities (“mixed” groups, the normal management used in previous hihi translocations). We ensured that mixing juveniles from different communities also included spatially-separated birds (i.e. only detected in northern or southern survey locations) that had little chance to interact prior to capture.

All birds for translocation were caught within 24 hours, then kept in the aviaries for four further days while samples were processed for disease screening. Each aviary held equal numbers of birds. During holding we provided supplementary food twice daily, using the same range of food used in previous successful hihi translocations (Ewen *et al*., 2018). On the evening of the 1^st^ April, hihi were re-caught from the aviaries, given standard health checks, and transferred to translocation boxes (five hihi per box). We transported all birds at the same time from the source site to the release site, overnight (by boat then van) to minimise stress for the birds. All hihi were released successfully the following morning (2^nd^ April).

### DEFINING FAMILIARITY AFTER TRANSLOCATION

We recorded associations at both the release site and source site from 3^rd^ April – 3^rd^ June 2017 in a similar manner as before translocation. However, as hihi were expected to disperse across the release site and not be fixed to locations following the translocation, we walked monitoring tracks at both sites to locate juveniles (by MM and CA at the release site: 300 observation hours, 38 individuals; by VF at the source site: 100 observation hours, 40 individuals). All three observers had similar experience of observing hihi as part of the standard monitoring of the source population. Whenever we encountered a juvenile, we noted the bird’s colour ring combination, the time it was first encountered (to nearest minute) and the time it left the area too quickly for us to follow. If we saw new individuals during the same time, we also noted their identity, entry time, and exit time. Using the same method at the release site and at the source site meant we could compare changes in social patterns in translocated juveniles to a group that had experienced network disruption while remaining in the same location. We constructed networks for source and release sites separately, using the same method as before translocation.

Post-release survival population surveys were conducted by MM at the release site every month between May – September 2017, and in March 2018 following the first breeding season of this new population. During each survey, MM walked monitoring tracks (a subset, alternated between surveys) across the release site for 40 hours over five days. Using this method meant birds could be detected by their calls across the entire site in each survey, located, and visually identified using binoculars. Partially-identified birds (for example, incomplete ring combinations) were discounted to limit misidentification. For each translocated hihi, we created encounter histories which represented each bird’s presence (“1”, seen) or absence (“0”, not seen) in each successive survey or “time point”. All individuals were assigned a “1” in time point 1 when they were released into the new site in April 2017 (all hihi were released successfully). Thus, an example encounter history would be “1110000”, where an individual was released at time point 1, seen in the surveys at time points 2 and 3, and then not seen again. Population surveys were also conducted at the source site by MM in September 2017 and February 2018 using the same method. For non-translocated hihi, we generated encounter histories from presence/absence in April 2017 observations, May 2017 observations, plus the two censuses. Time point 1 in these encounter histories (when all individuals were considered present and assigned “1”) was immediately before the translocation (March 2017). We used these data to investigate links between changes in social networks and survival for translocated birds, which we might expect if losing social connections was disruptive or stressful for hihi, and to compare their survival to non-translocated hihi.

### DATA ANALYSIS

#### Did translocation change group associations?

Social network analysis was conducted in R. We first tested if hihi grouped after the translocation according to familiarity based on (i) geographic distribution before translocation (north locations only, south locations only, mixed sightings); (ii) social network community before translocation (community 1–6); and (iii) aviary type during the translocation (“familiar” or “mixed”). For each analysis, we calculated the distribution of network edge weights and the assortativity coefficient (*r*, a value from +1 for total disassociation, to +1 for total association) to describe the strength of associations between juveniles based on our categorical measures (using the R package assortnet (version 0.12) (Farine, 2014)). We compared the *r* value of our network to the r values of 1000 random networks generated using pre-network data permutations in asnipe, to test if familiar juveniles were statistically more likely to associate than random. Data-stream permutations account for differences in the number of observations between individuals when calculating network statistics (Farine and Whitehead, 2015; Farine, 2017), and comparisons with permuted networks is a more robust method for determining statistical significance because networks are inherently non-independent and violate the assumptions of many statistical tests (Farine and Whitehead, 2015). Finally, we repeated our assortment analyses based on distribution and communities for the source site network after translocation, as a comparison from non-translocated birds. All *P*-values generated by comparing with permuted networks are specified as *P_rand_*.

#### Did individuals remain consistent in their sociality?

Second, we investigated if relative individual sociality remained consistent following translocation by comparing between translocated and non-translocated individuals. For each individual in each network, we calculated a weighted degree centrality (degree) which explained both its number and strength of associations. As the population sizes of juveniles were different before and after translocation, we then ranked individuals by their degree within each network, and divided ranks by the size of each population so all ranks were bound between 0 and 1. Thus, if individual sociality was consistent we would expect an individual’s rank to remain the same relative to others within their population. We assessed what predicted degree rank after translocation using a Generalised Linear Model (GLM) with a binomial distribution. Our predictors included degree rank before translocation, population (translocated or not translocated), and sex (translocations could affect male and female hihi differently (Armstrong *et al*., 2002, 2017)). We included an interaction between degree rank before translocation and population type, because sociality could be affected more extensively if moved to a new site. Finally, we also included number of observations after translocation as a fixed effect to ensure variation in degree rank was not only due to differences in detection among individuals.

To assess whether maintaining familiar groups during capture for translocations affected individual sociality, we calculated each translocated juvenile’s change in degree rank after translocation compared to before translocation (bound between –1 and 1; a negative value represented a decrease in social rank; a positive value was a rank gain). We used a Linear Model (LM) with rank change as the response. Our predictors included the aviary type each bird was housed in (“familiar” or “mixed” aviary) in interaction with degree before translocation (effects of aviary could depend on sociality), and sex. For this analysis, we included number of observations both before and after translocation as fixed effects, because change in rank score (our response) could be dependent on variation in both number of observations. Again, we assessed significance of both analyses using data-stream permutations.

#### Did social changes during the translocation affect survival?

Finally, we used our encounter histories for translocated birds to estimate survival depending on change in degree rank (−1 – 1: covariate) and sex (male or female: grouping factor) in Program MARK (version 9.0) (White and Burnham, 1999). As all individuals were identifiable, we used a live-recaptures (Cormack-Jolly Seber, CJS) analysis to estimate survival (*Φ*) and quantify re-sighting (*ρ*) to ensure survival was not confounded by varying re-sighting likelihoods between individuals. To ensure models explained variation in the data accurately, we first conducted a goodness-of-fit (GOF) test on a fully time- and group-dependent starting model, by calculating median ĉ as an estimate of overdispersion. We did not include covariates in this starting model as there is currently no method for GOF testing with covariates. The value for median ĉ = 1.30, which indicated a good fit of the data and so we corrected for the small level of overdispersion in further analyses. This meant we could accurately estimate *Φ* using:

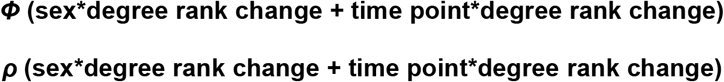

Here, we considered whether monthly survival was affected by the extent of change in rank degree after translocation compared to before translocation, explaining both loss and gain of associates relative to all other translocated juveniles. We considered rank change in interaction with sex, because this disruption could affect males and females differently, and time point (effects of social changes could vary across time). We accounted for variation in re-sighting likelihood with the same parameters.

We constructed a set of models with all combinations of predictors with and without the covariate, then ranked models by their corrected quasi-likelihood Akaike Information Criterion (QAICc, due to adjusting by median ĉ) values, which explain the model fit: a smaller QAICc value suggests the model better accounts for variation in the data. Any model less than 2 QAICc units from the top ranked-model was considered equally well supported. If multiple models had ΔQAICc < 2, we used model averaging to calculate effect sizes and 95% confidence intervals depending on model weight (which explained relative likelihood of each model). Any parameter with a confidence interval that did not span zero was considered to have a significant effect.

We analysed survival in non-translocated birds depending on degree rank change and sex in the same manner, to provide a comparison from birds remaining at the source site. However, we could not combine both translocated and non-translocated birds in one survival analysis to explore interactions with site statistically, as the time points of the surveys differed. Our median ĉ value following GOF was 1.42 for the starting source site model, which we corrected for in our analyses.

## Results

### DID TRANSLOCATION CHANGE GROUP ASSOCIATIONS?

Before the translocation, juvenile hihi formed robust communities which represented preferred and avoided associations (*r_comm_* = 0.71). Juveniles that were not translocated continued to be seen in the same areas of the island and with group-mates from the same communities before translocation (location: *r* = 0.12, *P_rand_* = 0.01, Table 1a, Figure 1; community: *r* = 0.14, *P_rand_* = 0.01, Table 1b, Figure 2). However, translocated juveniles behaved differently: they did not group according to either their geographic location (*r* = –0.04, *P_rand_* = 0.44, Table 2a, Figure 1) or community before translocation (*r* = –0.01, *P_rand_* = 0.19, Table 2b, Figure 2). Additionally, translocated juveniles did not associate more strongly if they had shared an aviary, even when they had been familiar at the source site; in fact, there was a tendency for a weak disassociation by aviary (Table 2c; *r* = –0.09, *P_rand_* = 0.04, Figure 2).

**Table 1.** Mixing matrices of association weights for hihi at the source site after the translocation based on (a) distribution before translocation at the source site (only in the “North”, “South”, or moved among the two “mixed”); and (b) network community before translocation (colours correspond to Figure 2a). 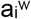 are the row sums, 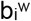 are the column sums; due to rounding, sum values may not be exact. Tables are symmetrical, so only half of values are shown.

| Distribution | North | Mixed | South | $a_i^w$ |
| --- | --- | --- | --- | --- |
| North | 0.21 | - | - | 0.41 |
| Mixed | 0.21 | 0.35 | - | 0.57 |
| South | 0.00 | 0.02 | 0.00 | 0.02 |
| $b_i^w$ | 0.41 | 0.57 | 0.02 | 1.00 |

(b)
| Community | red | yellow | blue | green | $a_i^w$ |
| --- | --- | --- | --- | --- | --- |
| red | 0.07 | - | - | - | 0.29 |
| yellow | 0.15 | 0.19 | - | - | 0.42 |
| blue | 0.04 | 0.03 | 0.11 | - | 0.19 |
| green | 0.03 | 0.04 | 0.00 | 0.03 | 0.10 |
| $b_i^w$ | 0.29 | 0.42 | 0.19 | 0.10 | 1.00 |

**Figure 1.**
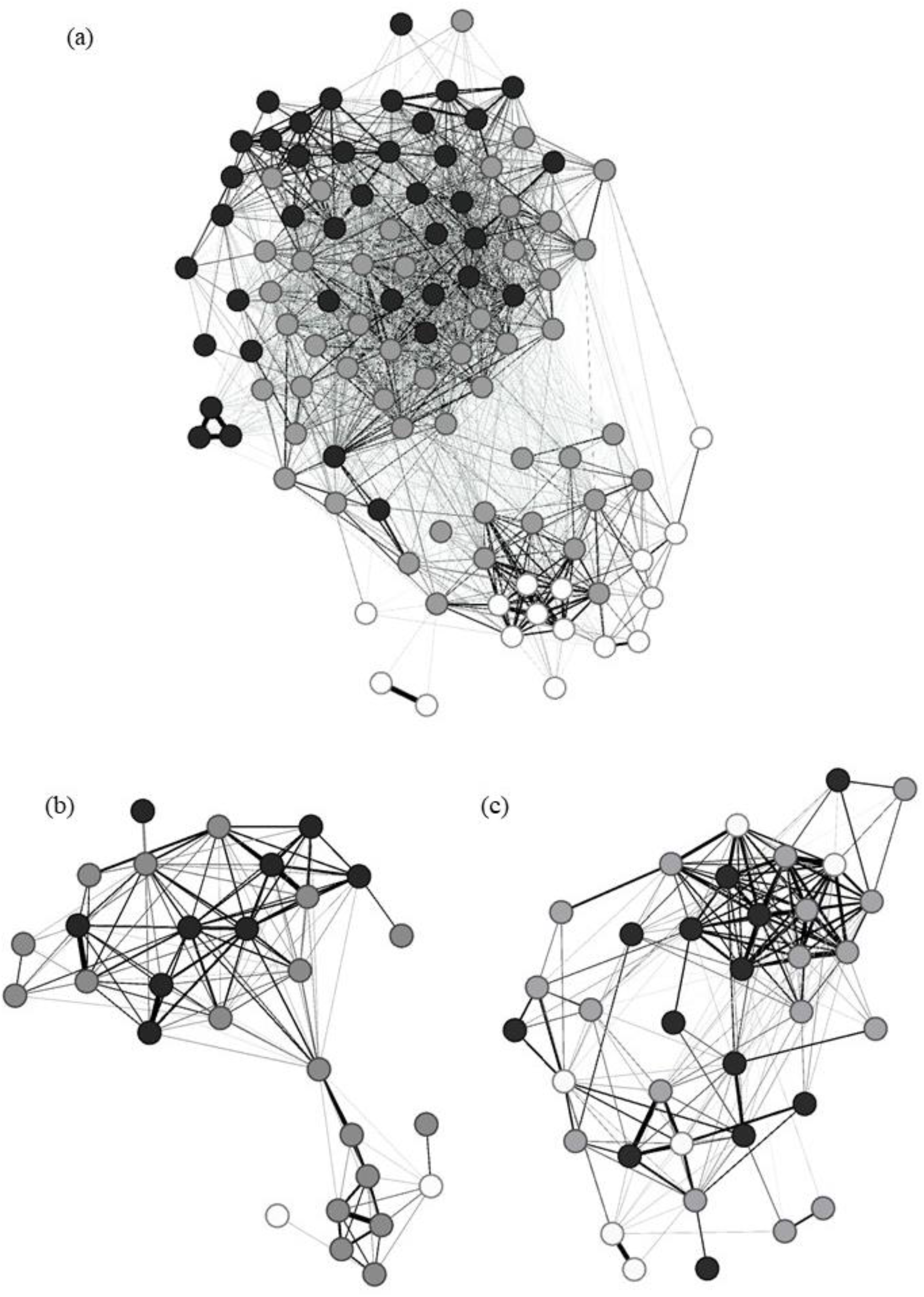
Hihi social networks (a) before translocation, and after translocation at (b) the source site, and (c) release site. Each node (circle) represents one hihi and the edges (lines) represent co-occurrence in a group. Edge width is proportional to association strength. Nodes in (a) are coloured by distribution at the source site (black = North; white = South; grey = mix). Nodes in (b) and (c) are coloured by the same distribution. Networks are arranged to minimise the length of edges between nodes which tends to cluster frequently-associating nodes together.

**Figure 2.**
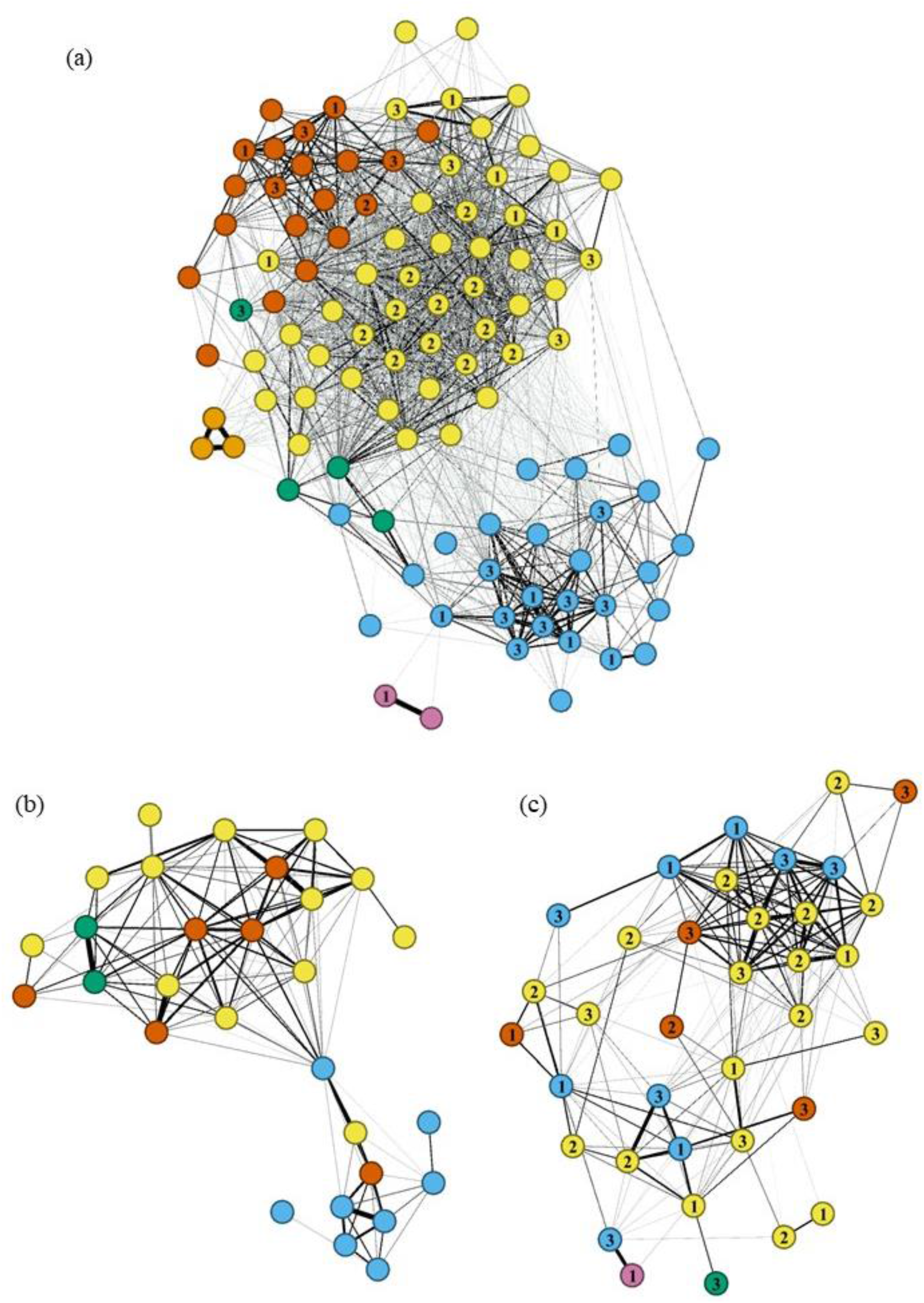
Hihi social networks (a) before translocation, and after translocation at (b) the source site and (c) the release site. Each node (circle) represents one hihi and the edges (lines) represents co-occurrence in a group. Edge width is proportional to association strength. Nodes in (a) are coloured by network community, and nodes in (b) and (c) are coloured by the same communities. Numbers in (a) and (c) correspond to the aviary each translocated juvenile was allocated. Networks are arranged to minimise the length of edges between nodes which tends to cluster frequently-associating nodes together.

**Table 2.**
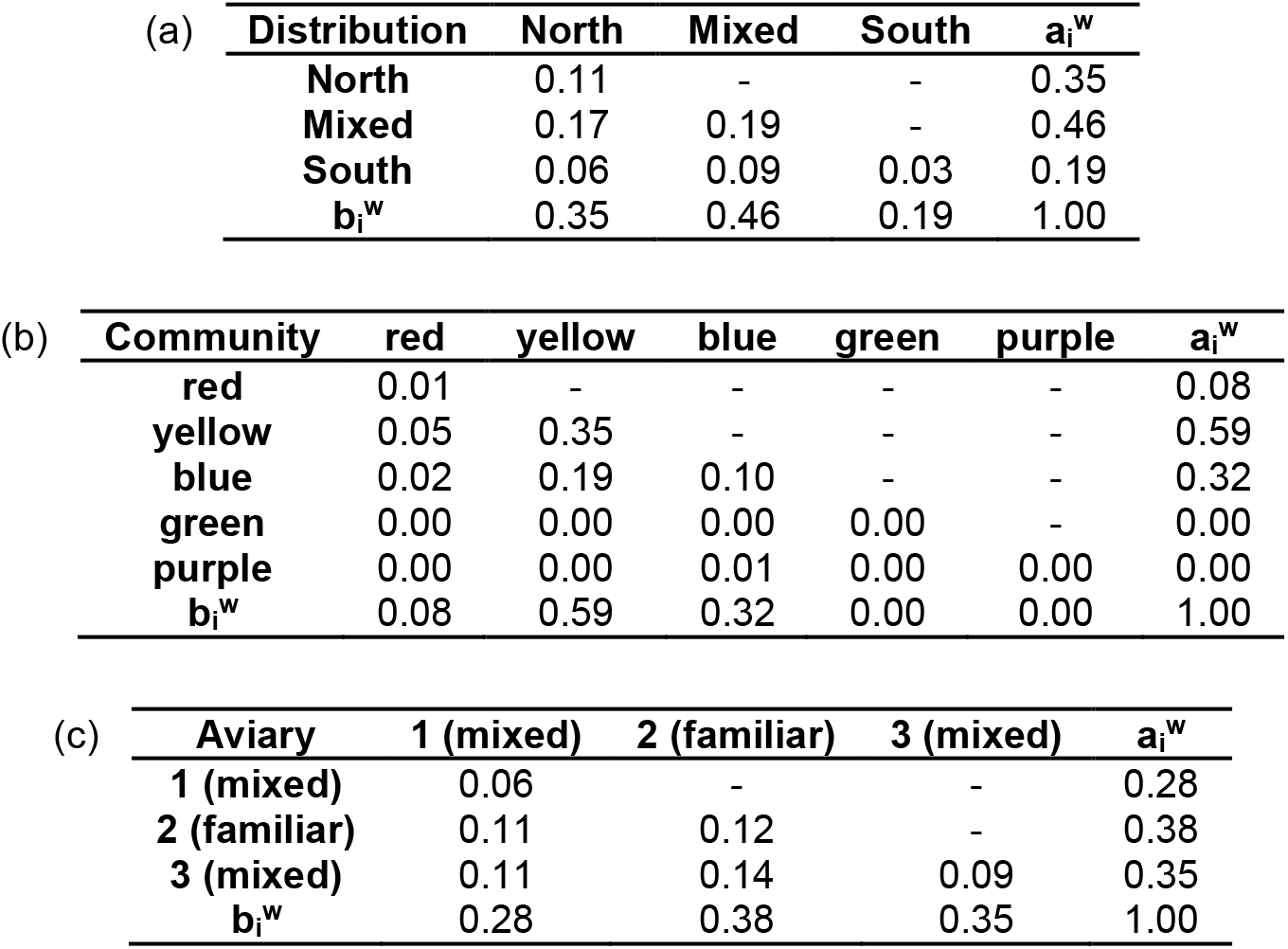
Mixing matrices showing association weights for hihi at the release site after translocation based on (a) distribution at the source site before translocation; (b) network community before translocation (colours correspond to Figure 2a); and (c) aviary number and category during translocation. 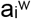 are the row sums, 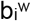 are the column sums; due to rounding, sum values may not be exact. Tables are symmetrical, hence only half of values are shown.

### DID INDIVIDUALS REMAIN CONSISTENT IN THEIR SOCIALITY?

Individual sociality was not consistent: more social juvenile hihi before translocation were not more social after the translocation at either the source site or release site (Table 3a, Figure 3a). Post-translocation social ranks did not differ between males and females (Table 3a) and also did not vary depending on how many times a bird was re-sighted any more than expected by random chance (Table 3a). Among translocated hihi, some birds experienced greater degree rank changes than others (greatest rank gain = +0.59; greatest rank loss = –0.68) but this was not predicted by their degree rank before translocation (both more- and less-sociable individuals were equally likely to change rank; Table 3b, Figure 3b). Individual degree rank was not preserved by holding a juvenile with its familiar group-mates in an aviary during the translocation (no significant difference in degree rank change between birds housed in familiar and mixed aviaries; Table 3b, Figure 3b). Finally, the extent of rank change was not significantly different between males and females (Table 3b), and again was not significantly affected by re-sighting before or after translocation compared to permuted networks (Table 3b).

**Table 3.** Results of (a) GLM analysing variation in post-translocation degree ranks and (b) LM analysing change in relative degree ranks for translocated hihi. Coefficients, standard errors and *z* or *t* values are presented. *P*-values generated from the original model are presented, but only for comparison to the *P*-values generated in relation to coefficients from 1000 randomised networks (*P_rand_*). Significant *P*-values are indicated in bold font.

| (a) | | coeff. | S.E. | $z$ | $P$-value | $P_{\text{rand}}$ |
| --- | --- | --- | --- | --- | --- | --- |
| degree rank after translocation ~ | intercept | -2.48 | 1.22 | -2.02 | **0.04** | 0.12 |
|  | before translocation degree rank | 1.19 | 1.53 | 0.78 | 0.44 | 0.24 |
|  | site (source site) | 2.10 | 1.35 | 1.56 | 0.12 | 0.16 |
|  | sex (male) | 0.29 | 0.52 | 0.55 | 0.59 | 0.13 |
|  | number of sightings after translocation | 0.18 | 0.07 | 2.59 | **0.009** | 0.27 |
|  | before translocation degree rank*site (source site) | -1.67 | 1.95 | -0.86 | 0.39 | 0.17 |
| (b) | | coeff. | S.E. | $t$ | $P$-value | $P_{\text{rand}}$ |
| change in degree rank (translocated hihi) ~ | intercept | -0.05 | 0.64 | -0.07 | 0.95 | 0.95 |
|  | degree rank before translocation | -0.84 | 0.80 | -1.05 | 0.30 | 0.75 |
|  | aviary category (mixed) | -0.04 | 0.66 | -0.06 | 0.96 | 0.69 |
|  | sex (male) | -0.05 | 0.06 | -0.90 | 0.37 | 0.22 |
|  | number of sightings after translocation | 0.34 | 0.01 | 8.26 | **<0.001** | 0.23 |
|  | number of sightings before translocation | 0.01 | 0.01 | 0.88 | 0.39 | 0.16 |
|  | degree rank before translocation*aviary category (mixed) | 0.20 | 0.72 | 0.27 | 0.79 | 0.51 |

**Figure 3.**
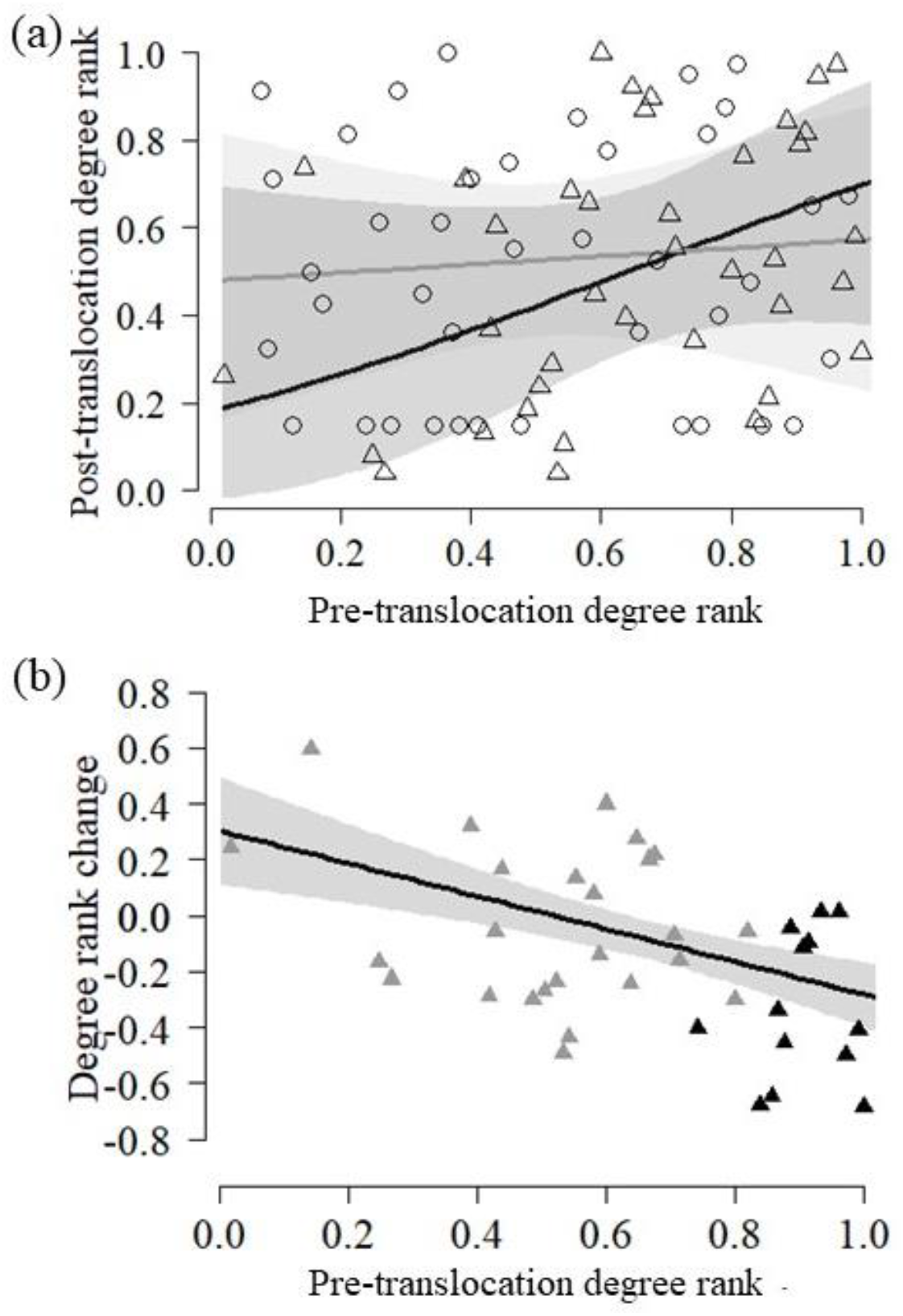
(a) relationship between degree ranks before and after translocation for non-translocated (circles, grey line) and translocated hihi (triangles, black line); (b) change in degree rank after compared to before translocation for translocated hihi held in mixed aviaries (grey triangles) and the familiar aviary (black triangles). Grey polygons represent 95% confidence intervals from models in Table 3.

### DID SOCIAL CHANGES ACROSS THE TRANSLOCATION AFFECT SURVIVAL?

Although we could not predict rank change, among translocated hihi there was a tendency for birds that experienced a greater decline in degree rank to have poorer post-release survival: the best supported model explaining monthly survival included rank change as a covariate, and sex, while accounting for varying re-sighting between sexes (Table 4; Supplementary Table 1a). However, monthly survival was high overall (Table 4) so the effects of degree change and sex were weak: models with no variation in survival were included in the set with ΔQAICc < 2 (Supplementary Table 1a). Survival rates were not time-dependent (Supplementary Table 5.1a), so we calculated overall 11-month survival likelihood based on monthly survival estimates from the models. 11-month survival showed greater variation, from 17.4% (95% CI = 0.2 – 66.7%) with the greatest loss of rank (−0.68) to 38.2% (95% CI = 3.0 – 78.4%) for the greatest rank gain (+0.59) (Figure 4). Overall male survival was 38.1% (95% CI = 12.7 – 64.6%) and female survival was 24.5% (95% CI = 3.3 – 57.5%) (Figure 4). For comparison, there was no evidence that degree rank change explained survival for non-translocated juveniles as it was included in models with little support (Supplementary Table 1b). In general, there was little support that survival varied with any predictor as many models were similarly ranked by ΔQAICc (Supplementary Table 1b).

**Table 4.** Initial model estimates of monthly post-release survival and re-sighting for translocated male and female juvenile hihi. Calculated from model averaging top-ranked models in Supplementary Table 1a.

| | Survival $\phi$ | | Re-sighting $\rho$ | |
| --- | --- | --- | --- | --- |
| | Effect ± SE | 95% CI | Effect ± SE | 95% CI |
| **Male** | 0.91 ± 0.03 | 0.80 – 0.96 | 0.94 ± 0.04 | 0.82 – 0.98 |
| **Female** | 0.88 ± 0.05 | 0.73 – 0.96 | 0.84 ± 0.07 | 0.67 – 0.93 |

**Figure 4.**
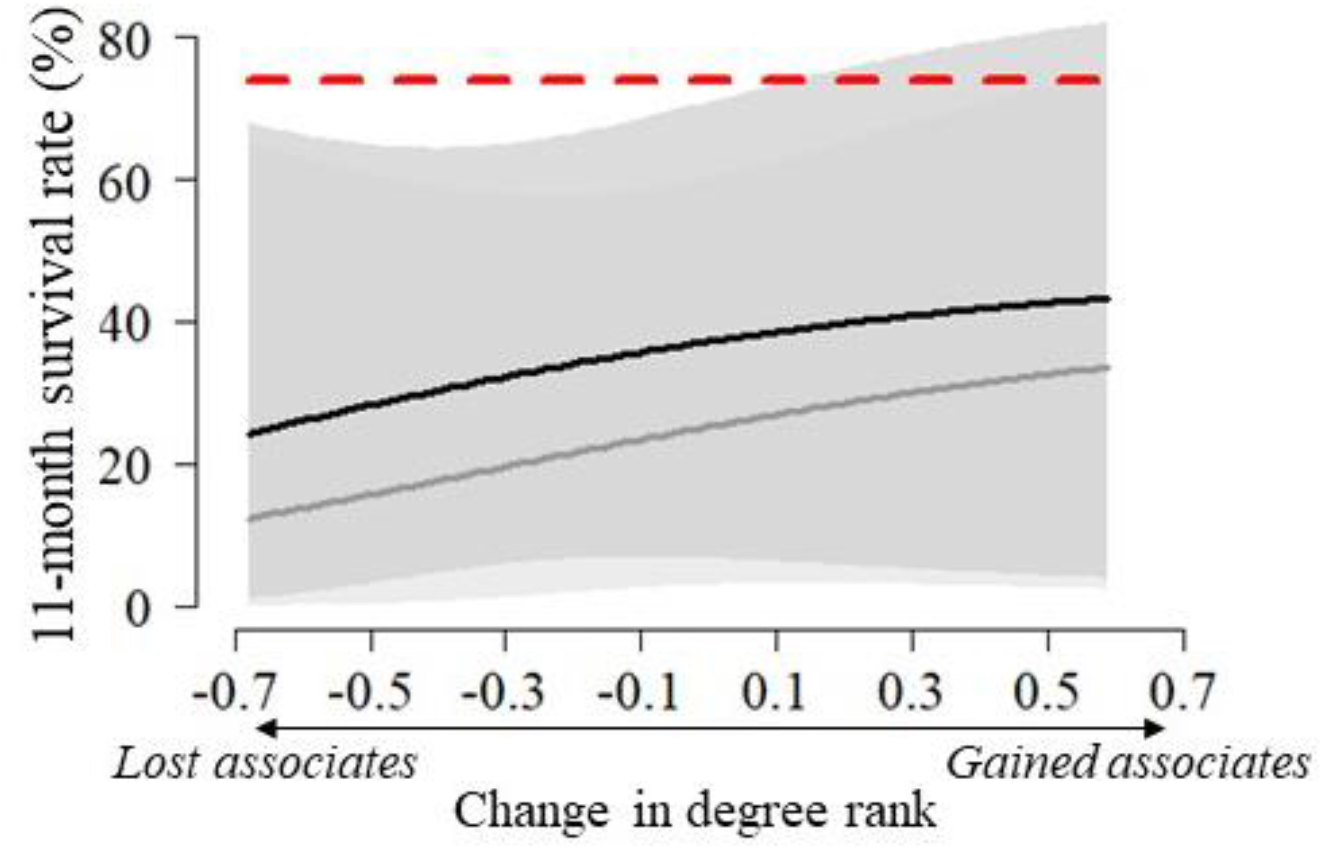
Predicted post-release survival likelihood across the 11 months post-release depending on change in degree rank after translocation, for males (black line) and females (grey line). 95% confidence intervals are grey polygons. Red dashed line represents survival estimate from non-translocated birds over the same length of time and shows no variation with degree rank change. All estimates predicted from model averaging top-ranked CJS survival models in Supplementary Table 1.

## Discussion

Here we have shown that translocating juvenile birds affects their social structure, which in turn may influence survival during the establishment phase of a reintroduction. Hihi that remained at the source site continued to associate with others from the same communities before translocation, but juveniles translocated to a new site formed new associations at random. Furthermore, holding juveniles together in an aviary did not promote group cohesion post-release, even if they had been previously familiar; instead, there was a suggestion that translocated birds actually disassociated from aviary-mates. At the individual level, there was no evidence that hihi maintained a similar level of sociality following a translocation event if they had been more social previously, and there was no difference between males and females; the same pattern was found in non-translocated hihi as well. Maintaining a group of familiar birds in an aviary did not prevent individuals from losing associates, relative to their previous sociality. Even though we did not find what predicted loss of sociality, translocated juveniles with the greatest decrease in their relative degree ranks showed a small, but significant, reduction in survival during their first year post-reintroduction. Our results suggest that translocation created a disruption to the social environment at both the group and individual level, and this may have consequences for likelihood of establishment.

Our finding that group structure changed during a reintroduction event, even when there was opportunity to maintain associations with familiar individuals (through translocating familiar hihi together), reflects results of earlier studies in other New Zealand bird species (Armstrong, 1995; Armstrong and Craig, 1995). These results were similar even though juvenile hihi are more social than the previous species studied and we also determined pre-translocation familiarity over a longer time period. Social disruption could be due to the process of translocating itself (catching, moving, and releasing) (Parker *et al*., 2012), which maintaining groups did not overcome. Alternatively, translocation could have removed external influences on associations when animals were removed from their original source environment. Understanding how environment and animal choice drive the formation of social groups is still in its infancy (Pinter-Wollman *et al*., 2013; Leu *et al*., 2016), but comparing with association patterns in a group of non-translocated birds meant we are able to draw stronger conclusions to suggest why social groups may not be maintained when moved to a new site. Only when hihi were removed from their source environment during the translocation did it result in mixing of previously less associated hihi; birds that experienced social disruption (removal of associates) while remaining in the source site maintained the same group structure. This suggests environment plays a key role in structuring hihi groups. Furthermore, it may mean that such groups will never be maintained during reintroductions, which by definition involve removing animals from one environment and placing them at a new site (Ewen, Armstrong, Parker, *et al*., 2012; IUCN/SSC, 2013).

Variation in number of associates between individuals influences a range of processes from how quickly animals find food (Aplin *et al*., 2012) to their risk of contracting disease (Christley *et al*., 2005). Associates may be particularly important when individuals need to rely on social information more: for example, when they have little personal information, such as following reintroduction to a new site (Kendal *et al*., 2005; Kendal, Coolen and Laland, 2009). In feral horses (*Equus caballus*), more social foals with a higher degree score were more likely to survive the loss of members of their herds during a “catastrophic” event that removed 40% of the population (Nuñez, Adelman and Rubenstein, 2015). Importantly, pre- and post-event sociality was not consistent for each foal, and post-event sociality was especially important for survival, which suggests the current social environment conferred the strongest advantages (Nuñez, Adelman and Rubenstein, 2015). In hihi, we found similar patterns as relative pre- and post-translocation sociality did not remain consistent for both translocated individuals, and birds that remained in the source environment (but did experience social disruption through the removal of peers). In our study, however, changes in sociality only had costs for survival when additionally associated with disruption of the abiotic environment. When translocated hihi lost more associates (and experienced the biggest disruption of their social environment) they showed a tendency to survive less well. We highlight that it may be this combination of disrupting both the social and physical environment that has the greatest consequences during the establishment phase of reintroductions. However, more work is needed to investigate why sociality changes; further data from translocations with lower survival may also help understand links between sociality and survival. Our release site was considered high quality for hihi (mature forest, assessed by expert members of the Hihi Recovery Group (Ewen, Adams and Renwick, 2013)) but conservation managers do not only use habitat quality to decide where to reintroduce, so future sites could be lower quality and have stronger survival pressures.

Holding animals together in temporary captivity pre-release is thought to promote group cohesion and improve the survival of translocated individuals in some species (Gusset, Slotow and Somers, 2006; Shier, 2006; Shier and Swaisgood, 2012; IUCN/SSC, 2013). However, we found the opposite direction of effect for hihi: birds kept in aviaries together showed a tendency for disassociation (suggesting avoidance) even if they had been familiar pre-capture. There was also no difference in degree rank changes between birds held in familiar and mixed groups. While all our familiar birds were ranked comparatively high for sociality, this was unlikely to be a confounder as they did not show any different trend compared to all other translocated birds. Therefore, in this species there does not appear to be a benefit of temporary captivity for maintaining or establishing a social environment. This complements previous research investigating other benefits of temporary captivity during hihi translocations. While temporary holding is a practical necessity due to the time needed to capture a required cohort, and is also used to reduce the risks of disease transmission (Ewen, Armstrong, Empson, *et al*., 2012), there is evidence that delaying release (even by four days instead of releasing immediately) decreases hihi post-release survival (Richardson *et al*., 2013). The downsides of captivity for some species such as hihi question how “soft” such delayed releases are (Batson, Abbott and Richardson, 2015), and highlights that there may be a need to tailor reintroduction protocols on a species-by-species basis, while considering multiple benefits and costs of management strategies (Moseby, Hill and Lavery, 2014).

Studies contrasting the effects of different treatments on conservation outcomes are essential to apply reintroduction biology effectively (Taylor *et al*., 2017). Our use of social network analysis provided a novel and detailed way to investigate the outcomes of conservation management for both group structure and individual sociality during a reintroduction (Snijders *et al*., 2017). By experimentally testing for changes in group structure and individual sociality during a reintroduction of hihi, our approach has provided important information for the management of this and similar species. In the case of hihi, translocation changed their group structure, which was further disrupted by holding groups in aviaries together. At the individual level, changes in associations (particularly loss of associates) was linked to mortality. Therefore, even if groups are not consistent, the quantity of associates may be important for juvenile survival during abrupt changes in the environment. Predicting this sociality change may need to be a focus of future work. Overall, we present one way that the importance of sociality can be tested, and highlight an as-yet little explored application for social network analysis to understand how social groups respond to our conservation interventions.

## Acknowledgements

We are very grateful to the Department of Conservation, members of the Supporters of Tiritiri Matangi, and staff and volunteers at Rotokare Scenic Reserve for their support during the translocation.

